# A functional genomic meta-analysis of clinical trials in systemic sclerosis: towards precision medicine and combination therapy

**DOI:** 10.1101/087361

**Authors:** Jaclyn N. Taroni, Viktor Martyanov, J. Matthew Mahoney, Michael L. Whitfield

## Abstract

Systemic sclerosis (SSc) is an orphan, systemic autoimmune disease with no FDA-approved treatments. Its heterogeneity and rarity often result in underpowered clinical trials making the analysis and interpretation of associated molecular data challenging. We performed a meta-analysis of gene expression data from skin biopsies of SSc patients treated with five therapies: mycophenolate mofetil (MMF), rituximab, abatacept, nilotinib, and fresolimumab. A common clinical improvement criterion of -20% OR -5 modified Rodnan Skin Score was applied to each study. We developed a machine learning approach that captured features beyond differential expression that was better at identifying targets of therapies than the differential expression alone. Regardless of treatment mechanism, abrogation of inflammatory pathways accompanied clinical improvement in multiple studies suggesting that high expression of immune-related genes indicates active and targetable disease. Our framework allowed us to compare different trials and ask if patients who failed one therapy would likely improve on a different therapy, based on changes in gene expression. Genes with high expression at baseline in fresolimumab non-improvers were downregulated in MMF improvers, suggesting that immunomodulatory or combination therapy may have benefitted these patients. This approach can be broadly applied to increase tissue-specificity and sensitivity of differential expression results.

## INTRODUCTION

Systemic sclerosis (SSc) is a rare systemic autoimmune disease characterized by skin fibrosis that lacks any FDA-approved therapies. To gain mechanistic insight into the action of experimental therapies, clinical trials in SSc have collected genome-wide gene expression data from skin biopsies pre- and post-treatment (Chakravarty et al., 2015; Gordon et al., 2015; Rice et al., 2015a; Hinchcliff et al., 2013; Chung et al., 2009)(Martyanov et al. Submitted). However, these studies in clinical trials face limitations. First, since SSc is a rare disease, clinical trials tend to have small sample sizes. Thus, few differentially expressed genes (DEGs) can be detected after multiple hypothesis testing correction (Chakravarty et al., 2015; Gordon et al., 2015). Second, not all therapy- or disease-relevant genes are regulated at the mRNA level. Identifying the functional consequences of treatments for SSc requires analytic methods beyond DEG analysis to infer the full biological impact of a therapy’s action.

In this study, we developed a machine learning method that uses “functional genomic networks” to learn the connectivity patterns of DEGs and extrapolate to the larger functional network in which a therapy is acting. We used our method to perform comprehensive analyses of five independent therapeutic trials in SSc, each with gene expression data. We complement our framework with Gene Set Enrichment Analysis (GSEA) (Subramanian et al., 2005) to identify differentially expressed pathways and find broad concordance with our network results.

Functional genomic networks are gene-gene interaction networks whose links encode functional relationships between genes, e.g., membership in the same pathway. These publicly available, tissue-specific networks were constructed using curated biological information (i.e., process annotations), as well as raw biological data (Greene et al., 2015). In our meta-analyses, we used linear support vector machine (SVM) classifiers to identify the connectivity patterns of the genes modulated during clinically significant treatment response. These patterns were then used to identify relevant non-DEGs. We show that this extrapolated set of genes includes the known therapeutic targets, even when these targets are not DEGs, demonstrating the power of our approach.

By examining multiple studies in parallel, we can identify the pathways commonly changed with clinical improvement, regardless of perturbation. We provide a comprehensive description of pathways altered during different treatments and identify molecular signatures characteristic of significant improvement. We show that non-responders from one trial (fresolimumab) have signatures that suggest possible response to other therapies. These results may be used to guide drug-repositioning efforts for patients that do not respond to a given treatment in clinical trials and ultimately for precision medicine in SSc.

## RESULTS

We analyzed publicly available gene expression data from clinical trials of five different therapeutic trials in SSc patients. These included the immunomodulators abatacept (Chakravarty et al., 2015) (CTLA4-IgG), mycophenolate mofetil (MMF) (Hinchcliff et al., 2013; Mahoney et al., 2015), and rituximab (Lafyatis et al., 2009; Pendergrass et al., 2012) (anti-CD20), a tyrosine kinase inhibitor (TKI) nilotinib (Gordon et al., 2015), and fresolimumab, targeting all isoforms of TGF-β (Rice et al., 2015a). All of these trials used skin disease severity measured by modified Rodnan Skin Score (mRSS) as one of the outcomes. We applied a common improvement criterion, which was a minimum decrease in mRSS of 20% *OR* 5 points (Khanna et al., 2006) post-treatment. The proportion of patients with available skin biopsy gene expression data who improved ranged from 27% (rituximab trial) to 83% (abatacept trial) (Table 1). The goal of this study was to understand the molecular processes perturbed by these diverse treatments, those that remain unchanged during each treatment, and to identify the differences between the subjects that did or did not improve.

**Table 1.**
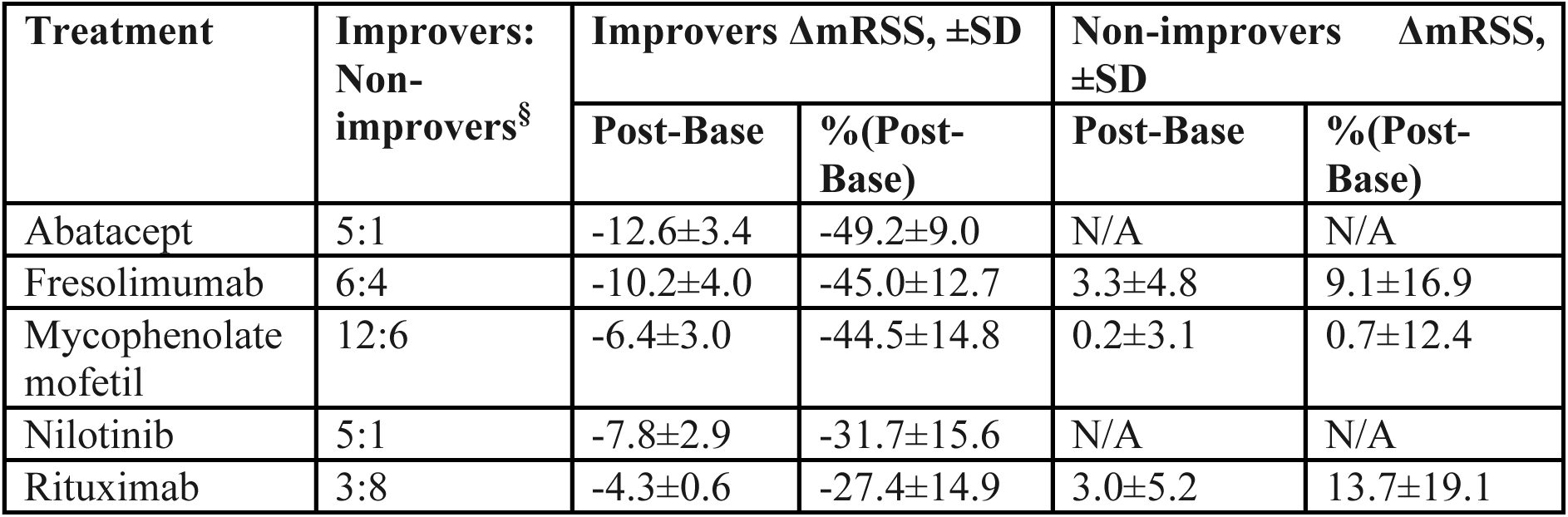
Overview of improvers and non-improvers based on the defined common improvement criterion. The absolute and percent changes in mRSS (± standard deviation) are reported. ^§^Only subjects with gene expression data were included.

### Network-based machine learning captures known treatment targets despite lack of differential expression

We developed a machine learning approach to identify pathways down-regulated by treatment (Figure 1). Genes with nominally significant decrease post-treatment (uncorrected p <0.05, paired t-test) in clinically significant improvers were supplied as positive examples to a SVM classifier (Greene et al., 2015)(Figure 1); genes that showed no evidence of differential expression are used as negative examples (0.95<uncorrected p≤1). The classifier learned the connectivity patterns of the DEGs in the Genome-scale Integrated Analysis of gene Networks in Tissues (GIANT) skin network and returned a ranking of all genes in the genome (Greene et al., 2015). Genes with high positive scores are most functionally similar to the nominally significant DEGs from the expression analysis (Greene et al., 2015), but are not required to be differentially expressed. Thus, top-ranked genes may be unregulated at the mRNA level or ‘missed’ due to small sample sizes, but are highly relevant to response. Similar approaches have been applied to GWAS re-analysis (Greene et al., 2015) and to DEGs for novel viral antagonism mechanism detection (Gorenshteyn et al., 2015).

**Figure 1.**
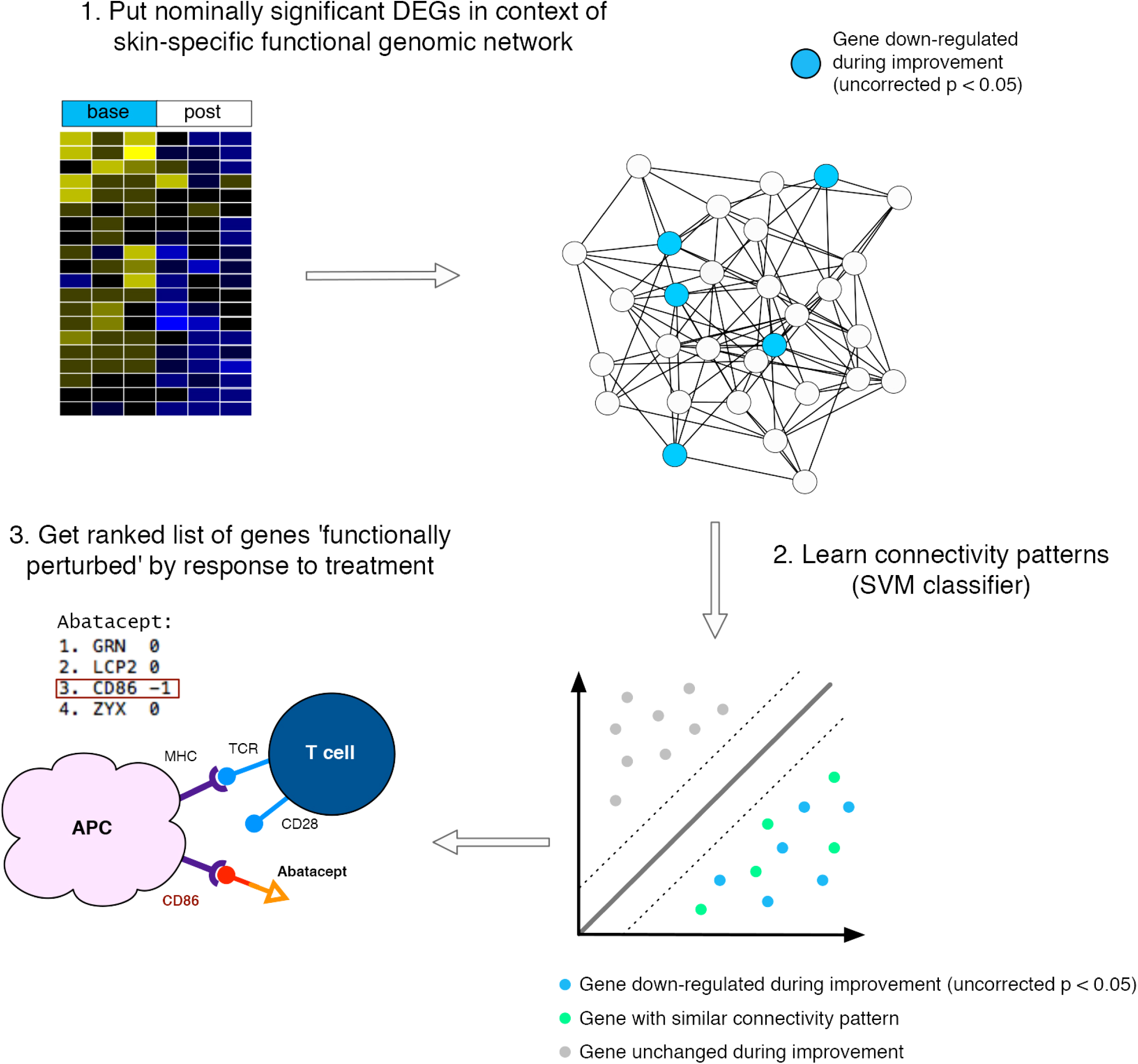
Schematic overview of machine learning approach. Nominally significant genes that decrease post-treatment (uncorrected p < 0.05) in clinically significant improvers are supplied as positive examples to a linear SVM classifier (DEGS); genes that show no evidence of differential expression are used as negative examples. The classifier learns the connectivity patterns of the DEGS in the GIANT skin network and returns a ranked list of gene symbols (highly positive: most like nominally significant DEGS). In the case of abatacept, the classifier returns *CD86* as a highly ranked gene — one of the molecules that interacts with this biologic — despite being a negative example.

As a positive control, we tested whether our approach could prioritize known drug targets better than differential expression alone. Abatacept is a cytotoxic T-lymphocyte–associated antigen 4–immunoglobulin fusion protein (CTLA-4–Ig) that binds to CD80 or CD86. Using data from an investigator-initiated trial of abatacept in SSc patients (Chakravarty et al., 2015), the classifier returned *CD86* as the third highest-ranked gene, despite not being differentially expressed (p=1) (Figure 1). Differential expression analysis missed this highly relevant gene, but the DEGs were functionally related to *CD86* enabling our approach to find it. Beyond single gene targets, we found that our method is better at capturing relevant target gene sets than the t-statistic alone (Figure 2a and 2b), which is used in many DEG approaches.

**Figure 2.**
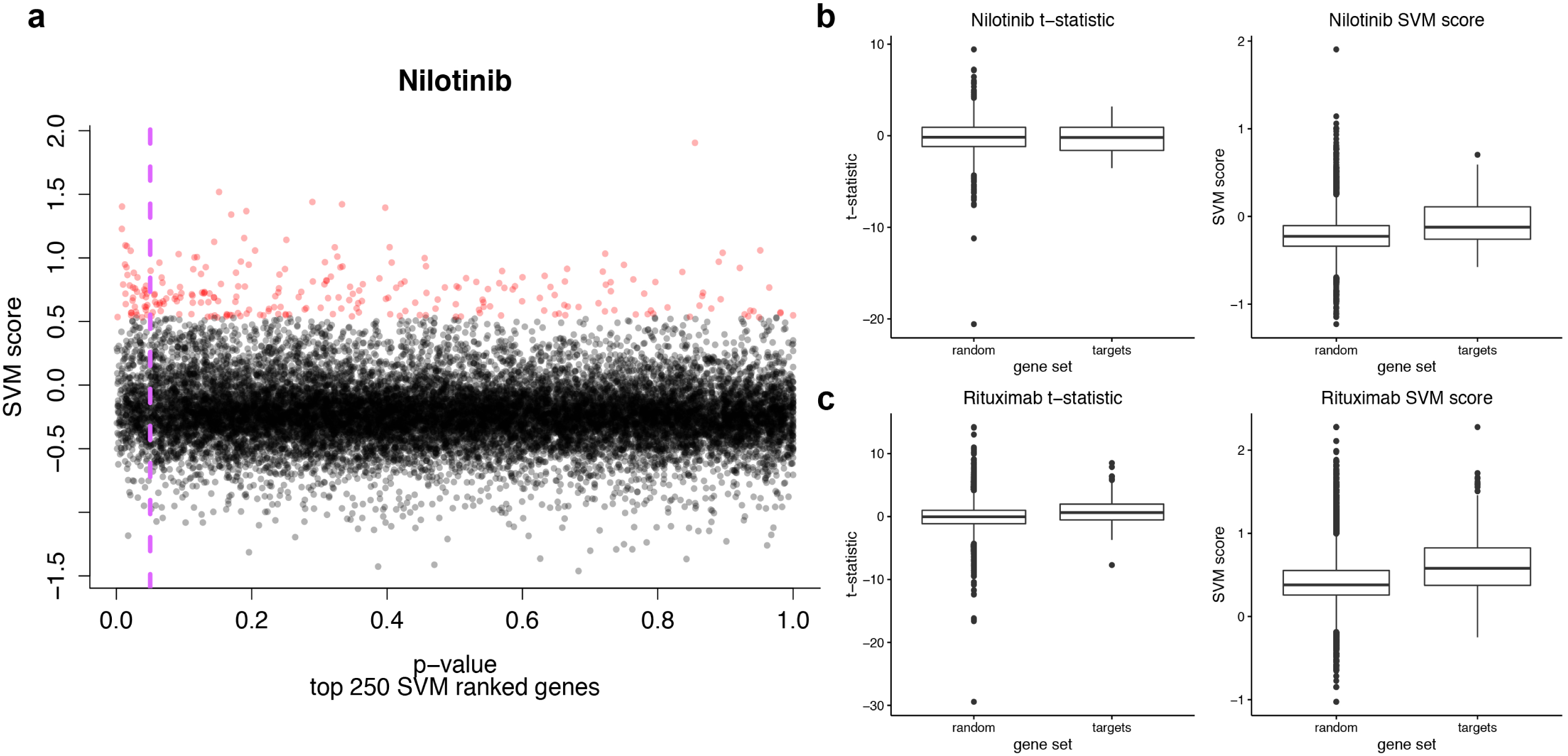
The linear SVM classifier captures features beyond just differential expression and better separates nilotinib targets from random sets of genes than the t-statistic alone. (a) Scatterplot of SVM scores vs. p-values. The genes with the highest SVM scores are highlighted in red. The purple dashed line indicates a p-value=0.05. (b) Boxplots of SVM scores and t-statistics comparing the target gene set vs. one hundred gene sets of the same size (“random”). There is no significant difference in the t-statistic distributions (Mann-Whitney-Wilcoxon p=0.73), whereas the difference in the SVM scores distributions is significant (Mann-Whitney-Wilcoxon p=0.0004). (c) B cell gene distributions (as annotated at the protein level using the Human Protein Reference Database) in the rituximab data - t-statistic Mann-Whitney-Wilcoxon p=1.75×10^-9^, SVM score Mann-Whitney-Wilcoxon p<2.2×10^-16^.

Nilotinib is a TKI with clear sets of known target genes (kinases; taken from Yoo et al. (2015) [D2]). These targets show higher SVM scores than random gene sets of the same size (p=0.0004), but there is no significant difference in t-statistics between target and random genes (p=0.73) (Figure 2b). Our approach also sheds light on how cell types are perturbed during treatment. Rituximab has been shown to deplete dermal B cells (Lafyatis et al., 2009). B cell-specific genes, as determined by the Immune Response in silico study (IRIS; p=0.029) (Abbas et al., 2005) and the Human Protein Reference Database (HPRD; p<2.2×10^-16^) (Prasad et al., 2009) have higher SVM scores than random. In contrast, t-statistics of B cell genes are either no different than random (IRIS p=0.38) or the difference is less statistically significant (HPRD p=1.75×10^-9^) (**Figure S1 and 2c**).

### Improvement-associated gene signatures indicate common pathways of skin disease resolution

We then investigated if the SVM-generated gene signatures for each therapy were functionally associated with one another using a z-score method (Huttenhower et al., 2009) (**Figure S2a-c**). This approach quantifies whether the top-ranked genes (250) from the SVM are more strongly connected to one another in the skin functional network than random sets of genes of the same size. If two gene sets have a z-score>3, that indicates that they are significantly connected and therefore are inferred to participate in similar pathways. Every pair of gene signatures from each of the five trials was highly significant (**Figure S2a-c**). Notably, improver signatures are generally more significant than non-improver signatures (**Figure S2b**) or treatment-effect alone (**Figure S2c**), suggesting that there is a core biology underlying the resolution of SSc skin disease.

### Complex network theory identifies skin-specific functional gene sets

To detect the common pathways underlying skin disease resolution, we used community detection to identify functional modules in the GIANT skin network (Greene et al., 2015) (see Methods). Because of the way these networks were constructed, community detection identifies sets of genes that participate in coherent biological processes in a *tissue-specific* manner. We determined which functional modules had high or low SVM scores to ascertain what pathways were down-regulated or unchanged by treatment, respectively. Figure 3 illustrates functional modules that have significantly high or low SVM scores as compared to the entire distribution (Wilcoxon test, Bonferroni-adjusted p<0.001). Below, we report functional modules that had high SVM scores for multiple therapies, as these are indicative of ‘overlapping’ modulated biology in multiple trials (we restricted further study to the top 20 modules in each trial for brevity). Generally, bottom-ranking functional modules encoded housekeeping processes (e.g. ribosome biogenesis and mRNA processing). This is a useful positive control because we do not expect therapies to target such processes.

**Figure 3.**
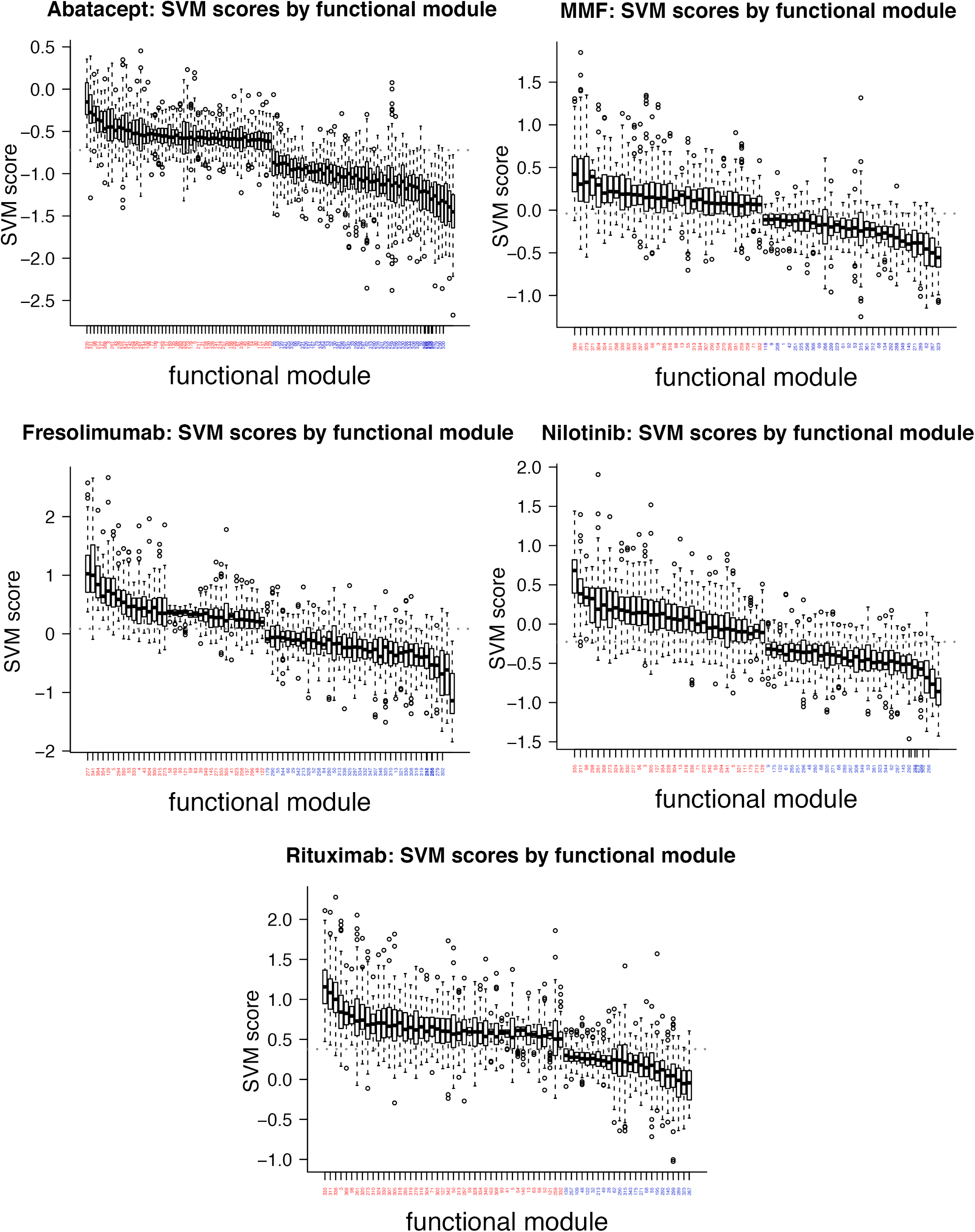
Boxplots of functional modules with significantly high or low SVM scores (Bonferroni adj. p<0.001). A red label indicates high SVM scores; blue indicates low scores. The gray dashed line indicates the median SVM score in each case.

A module (273) enriched for fibrosis-related processes such as *response to TGF-β* and *signaling by platelet-derived growth factor (PDGF)* was highly ranked in all studies but abatacept (Table 2). Modules enriched in immune-related processes were highly ranked in all studies except fresolimumab Table 2. This suggests that fresolimumab, a monoclonal antibody to TGF-β, does not alter the same immune-related processes as other therapeutics, consistent with its mechanism of action. Additionally, the perturbation of a module enriched for Interleukin-6 signaling (261), which is specifically targeted by tocilizumab (anti-IL-6) (faSScinate trial) (Khanna et al., 2015), highlights the central importance of IL-6 in SSc skin disease.

**Table 2.**
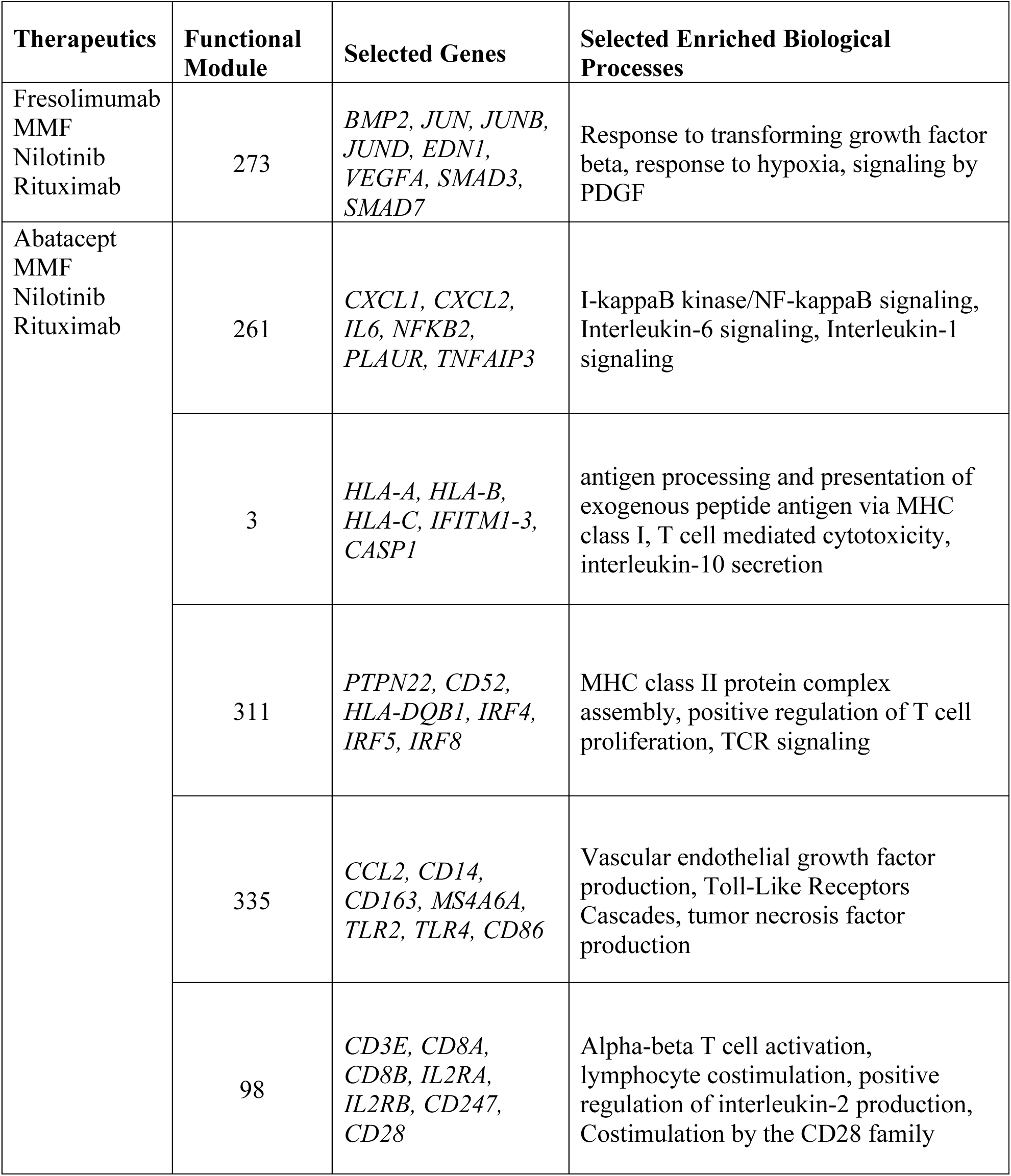
Functional modules predicted to be down-regulated in four out of five therapeutic trials. Top 20 significant modules only. Selected member genes and biological processes are shown. Functional enrichment was performed using gProfileR (Reimand et al., 2011).

### Different immunomodulatory treatments modulate distinct functional processes

Intriguingly, no top modules were shared between abatacept and MMF *only* (modules in common across abatacept, MMF, and at least one other treatment were found, Table 2). This is despite the fact that improvers in both studies have been reported to have high ‘inflammatory’ signatures pre-treatment that were down-regulated post-treatment (Hinchcliff et al., 2013; Chakravarty et al., 2015). The lack of additional overlapping modules suggests that there may be heterogeneity within the inflammatory intrinsic subset (Milano et al., 2008; Pendergrass et al., 2012; Hinchcliff et al., 2013) that can be targeted by one or the other therapy. To contrast the functional targets of these two therapies, we standardized their SVM scores in order to make a direct numerical comparison and identified those functional modules that significantly differed between them (**Figure S3**). This showed that T lymphocyte-, vascular-, and proliferation-related gene sets are likely to be differentially affected by abatacept and MMF (**Table S1**). Abatacept had higher standardized scores for vascular- and collagen-related modules (129 and 277), although it is possible that this is due to the greater magnitude of the improvement in the abatacept trial (Table 1). MMF had higher scores for proliferation (module 285 and 302) and type I interferon modules (module 336). This may be due to the broadly immunosuppressive nature of MMF, which suppresses lymphocyte proliferation, compared to the molecularly precise targeting of abatacept, which inhibits T cell co-stimulation of antigen presenting cells. Overall, there were fewer genes with positive SVM scores for abatacept.

### Comparison of network analyses with GSEA results

As a final control for our analyses, we used GSEA to test each study for differential expression of sets of genes (pathways) rather than single genes. GSEA is a method for identifying gene sets that are altered between two phenotypes or timepoints. It is a well-established procedure that is complementary and provides further validation to our network approach. We used a collection of curated “Hallmark” gene sets (Hallmarks) that “summarize and represent specific well-defined biological states or processes and display coherent expression” (Liberzon et al., 2015). The major limitation of GSEA compared to our approach is the requirement that users identify relevant gene sets (in this case Hallmarks) *a priori*. Nevertheless, the GSEA results were broadly concordant with our network results.

We focused on processes in common between studies, which represent biological processes relevant to disease resolution either due to intervention or spontaneous improvement. A single Hallmark, *epithelial-mesenchymal transition* (EMT), was significantly decreased post-treatment in improvers from all five studies. All studies except for fresolimumab modulated Hallmarks involved in the immune system signaling, e.g. *IL6/JAK/STAT3 signaling* and *TNFA/NFKB signaling* (**Table S2**). This agrees with our network-based functional module results (Table 2; modules 261, 3, 311, 335, 98), where we found that immune-related processes were down-regulated by all therapies except for fresolimumab.

Abatacept, fresolimumab, nilotinib and rituximab all resulted in changes in the *TGFB signaling* Hallmark (**Table S2**), but no change in this pathway was observed with MMF treatment. This is somewhat in contrast with our network results, where we found that all therapeutics but abatacept down-regulated genes implicated in *response to transforming growth factor beta* (found in module 273). Some differences between the two methods are to be expected. The identification of functional modules is data-driven, in contrast to the expert-curated Hallmarks, and we restricted our analysis to top modules only, not all that reached significance. However, there were two common “core enrichment genes” that contributed to the significant enrichment score for the *TGFB signaling* Hallmark in all four studies: *THBS1* and *SERPINE1* (**Table S2**). Both are strongly correlated with mRSS (Farina et al., 2010; Rice et al., 2015b). These genes are in module 304, which is one of the top 20 modules for MMF. This suggests that while MMF did not significantly down-regulate the *TGFB signaling* Hallmark by GSEA criteria, it is hitting a functionally similar set of genes by our networks method. Likewise, although abatacept did not down-regulate genes associated with *TGFB signaling*, it did down-regulate genes enriched for the Reactome pathway *Degradation of the extracellular matrix* (module 277). Thus, although the pathways that significantly overlap between studies depend somewhat on the method used, the down-regulation of collagen or extracellular matrix deposition pathways is commonly found in all studies. In addition, both methods identified differences between the immunomodulatory treatments abatacept and MMF, further suggesting that these two therapies may resolve skin disease severity differently. Most importantly, fresolimumab is the only therapy that does not appear to alter immune-related processes as determined by either method.

### Fresolimumab non-improvers may benefit from immunomodulatory treatment

Original publications of these studies largely ignored the molecular changes measured in non-improvers. However, our results suggest that similar processes were down-regulated in non-improvers post-treatment across multiple therapies (**Figure S2b**). We hypothesized that the use of non-improver signatures could help distinguish between processes that were truly down-regulated due to improvement, compared to those that were affected due to treatment alone, and would allow us to identify therapies that may be more clinically effective for a particular patient.

We used an SVM classifier on the GIANT skin network to classify genes that were uniquely down-regulated in improvers post-treatment (uncorrected p<0.05); negative examples were genes uniquely down-regulated in non-improvers. This resulted in genes with highly positive SVM scores being most like genes down-regulated in improvers and genes with highly negative SVM scores being most like genes down-regulated in non-improvers. We refer to these as therapeutic ‘post lists’ below. We performed a similar analysis to identify genes most like those elevated in improvers (highly positive scores) or in non-improvers (highly negative scores) pre-treatment (termed ‘base lists’; see Methods).

We determined if subjects that failed to respond to fresolimumab (non-improvers) may have had active inflammatory pathways that were modulated by one of the other therapies (i.e. identifying complementary treatments). Functional enrichment analyses for the bottom 250 genes in the fresolimumab base list (most like genes elevated in non-improvers pre-treatment) showed that they were significantly enriched for *lymphocyte aggregation* and *type I interferon production*. We then asked whether fresolimumab non-improvers were likely to have responded to one of the immunomodulatory treatments analyzed, such as MMF.

We determined if bottom-ranking genes in the fresolimumab base list were high-ranking genes in the MMF post list—that is, if genes most like those elevated in fresolimumab non-improvers were also genes most like those down-regulated uniquely in MMF improvers. First, we compared the fresolimumab base and MMF post lists using a metric called rank biased overlap (RBO; the extrapolated version (Webber et al., 2010)), which is 1 if they are exactly the same, 0 if they are exactly opposite, and 0.5 if the rankings are random with respect to each other (Figure 4a). The fresolimumab base and MMF post ranked lists have RBO=0.34, suggesting that they are significantly dissimilar. We also asked if the bottom 250 genes from the fresolimumab base list were enriched near the top of the MMF post list using a pattern-matching strategy (Lamb et al., 2006); the KS statistic was highly significant (permuted p<0.001) indicating that was the case. Finally, we showed that bottom-ranked fresolimumab base gene sets of various sizes had significantly highly positive MMF post SVM scores (Figure 4b). These results suggest that fresolimumab non-improvers not only had immune-related signatures active pre-treatment and not modulated by therapy, but also that these pathways may have been modulated by MMF treatment (Figure 4c).

**Figure 4.**
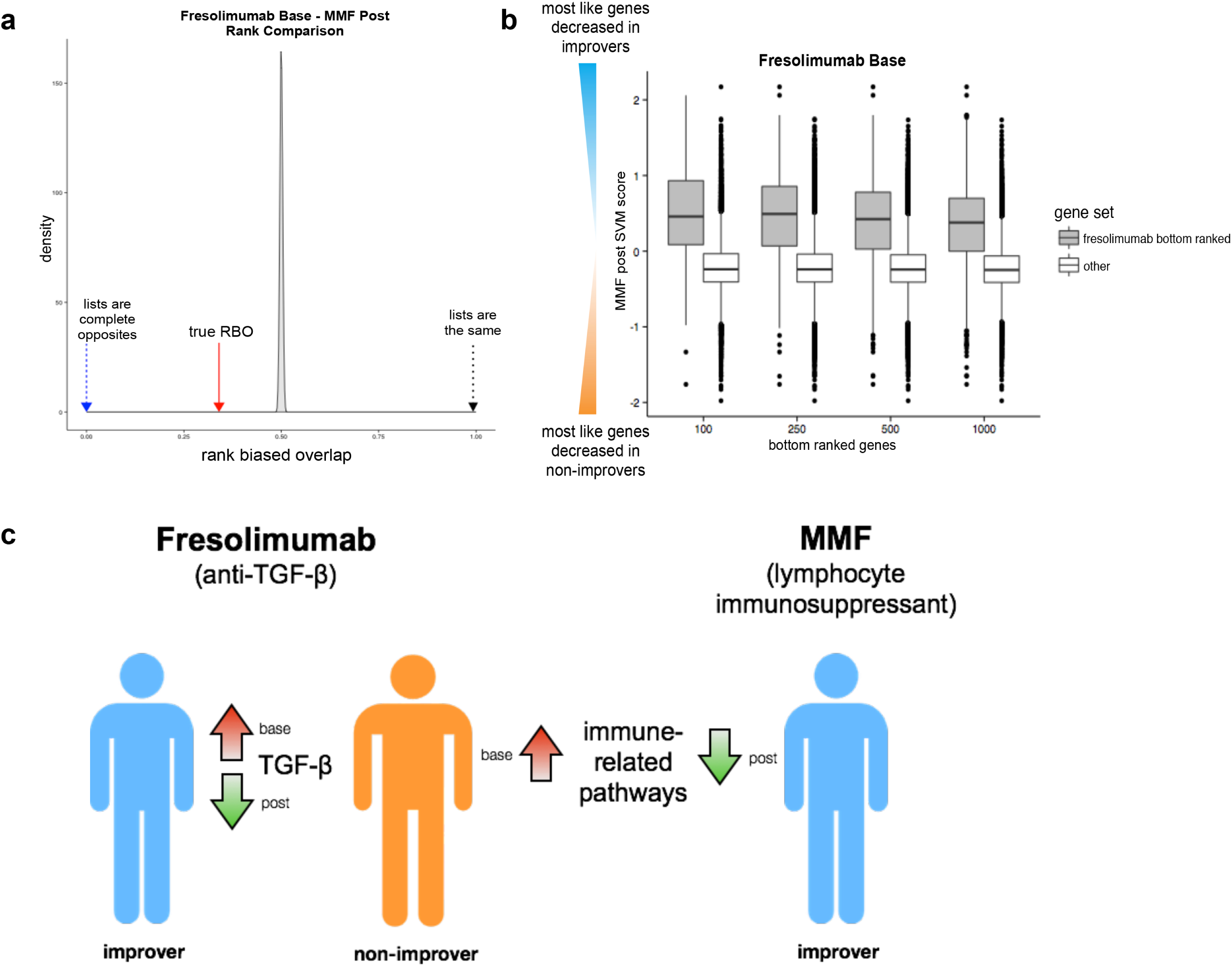
Several complementary methods suggest fresolimumab non-improvers may have benefitted from MMF treatment. (a) Density plot of the rank biased overlap (RBO) null distribution (permuted SVM scores). The red arrow indicates the true RBO (0.34) between fresolimumab base and MMF post ranked lists. (b) Boxplot illustrating that fresolimumab base bottom ranked gene sets of increasing size (100, 250, 500, and 1000 genes) have significantly highly positive MMF post SVM scores. (c) Schematic overview of findings. Fresolimumab non-improvers have elevated immune-related genes pretreatment; immune-related genes are uniquely decreased in MMF improvers.

## DISCUSSION

Successfully treating SSc manifestations requires modulation of many biological pathways, which must occur downstream of a therapeutic molecule binding to its small set of targets. Genome-wide gene expression data are now routinely gathered in clinical trials and provide insight into the functional consequence of treatment. A limitation is that most analyses only examine genes with statistically significant changes at the mRNA level, which may not fully capture the functional consequences of treatment.

We developed an approach that leverages functional genomic networks and machine learning to extrapolate from DEGs to the broader functional context in which a therapy is acting. Our strategy goes well beyond differential gene expression to identify the core processes that change in response to therapy and their critical component genes. As a positive control, we found that while a therapeutic’s targets are not always differentially expressed, the DEGs are highly functionally similar to the known therapy targets.

We cannot rule out the possibility that the subject composition of each of the cohorts included in this work influenced our results. However, we show that by putting nominally significant DEGs in the context of functional networks we can identify highly relevant genes (e.g., target kinases for nilotinib, B cell genes for rituximab) that the original studies did not identify as significant (Figure 2 and S2). Furthermore, our machine learning approach allows us to take advantage of the presence of non-improvers in some studies and apply conservative cutoffs for gene sets of interest.

Our results add to the growing body of evidence that molecular phenotyping in SSc patients prior to treatment may increase the likelihood of meaningful clinical response. In particular, we observe the abrogation of inflammatory pathways in multiple trials regardless of a therapy’s mechanism of action, which supports the hypothesis that the high expression of immune-related genes may represent an active disease state that is most clinically actionable.

Across multiple studies, improvers were characterized by post-treatment down-regulation of immune and fibrotic Hallmarks. Of particular interest was the modulation of *EMT* in improvers from all trials. Genes comprising the *EMT* Hallmark showed significant enrichment in *extracellular matrix organization* (ECM) and ECM-related functional terms e.g. *cell adhesion, vasculature development* and *collagen formation*. This corroborates our previous work showing that an ECM functional module occupies a central part in the putative network structure of SSc (Mahoney et al., 2015).

However, we also noted subtle differences in the pathways modulated by abatacept and MMF, both immunomodulatory therapies, suggesting that there is no therapy that is a ‘panacea’ for the inflammatory subset as of yet. Conversely, fresolimumab was shown to be less likely to down-regulate immune pathways than the other therapeutics and fresolimumab-treated non-improvers had elevated inflammatory processes at baseline. This suggests that if a baseline inflammatory signature is present in a patient they may benefit from an immunosuppressant, perhaps in combination with anti-TGF-β therapy.

In this work we addressed some of the special considerations of pilot studies in a rare disease, and we aimed to be conservative, particularly because multiple trials were under examination. As the field continues to conduct small trials, it will be important not only to amass more molecular data, but also to build new frameworks to analyze and interpret them. Our results show that functional genomic networks are a powerful complement to purely statistical techniques. By extrapolating from a noisy list of DEGs, we have identified common mechanisms of action for multiple therapies, as well as critical differences between them.

## MATERIALS & METHODS

### DEG analysis

DEG analysis was performed using the Comparative Marker Selection GenePattern module (Reich et al., 2006) using default parameters. In the case of pre- and post-treatment comparisons, paired t-tests were used; for all other comparisons, unpaired t-tests were used. The t-statistics and uncorrected p-values used throughout the text are taken from this analysis.

#### Functional genomic network analyses

All SVM classifiers were implemented using the Network-guided GWAS Analysis from the GIANT webserver (http://giant.princeton.edu) and using the GIANT skin network as the context (Greene et al., 2015). For improver signatures, the positive examples were genes down-regulated post-treatment in improvers (uncorrected p<0.05) and the negative examples were genes unchanged post-treatment (0.95<uncorrected p≤1). The same approach was used for non-improver signatures. To generate ‘base’ ranked lists, the positive examples were genes that were higher in improvers pre-treatment (uncorrected p<0.05) and the negative examples were genes that were higher in non-improvers pre-treatment (uncorrected p<0.05). To generate ‘post’ ranked lists, the positive examples were genes down-regulated in improvers post-treatment (uncorrected p<0.05) and the negative examples were genes down-regulated in non-improvers post-treatment (uncorrected p<0.05). For the boxplots in Figure 2b and 2c and density plots in **S1a** and **S1b**, 100 gene sets of the same size as the target or B cell gene sets were randomly sampled and one-sided Mann-Whitney-Wilcoxon tests were used to compare the distributions. More information on functional genomic and network methods is in the Supplemental Material.

## ACKNOWLEDGMENTS

JNT would like to thank members of the Whitfield Lab, J. K. Gordon, and C. S. Greene for helpful discussion.

